# A mechanistic model explains oscillatory slowing and neuronal hyperactivity in Alzheimer’s disease

**DOI:** 10.1101/2022.06.20.496731

**Authors:** Christoffer G. Alexandersen, Willem de Haan, Christian Bick, Alain Goriely

## Abstract

Alzheimer’s disease is the most common cause of dementia and is linked to the spreading of pathological amyloid-*β* and tau proteins throughout the brain. Recent studies have highlighted stark differences in how amyloid-*β* and tau affect neurons at the cellular scale. On a larger scale, Alzheimer’s patients are observed to undergo a period of early-stage neuronal hyperactivation followed by neurodegeneration and frequency-slowing of neuronal oscillations. Herein, we model the spreading of both amyloid-*β* and tau across a human connectome and investigate how the neuronal dynamics are affected by disease progression. By including the effects of both amyloid-*β* and tau pathology, we find that our model explains AD-related frequency slowing, early-stage hyperactivation, and late-stage hypoactivation. By testing different hypotheses, we show that hyperactivation and frequency-slowing are not due to the topological interactions between different regions but are mostly the result of local neurotoxicity induced by amyloid-*β* and tau protein.

## Introduction

Dementia is a syndrome characterized by gradually progressive memory dysfunction, language impairment, behavioral aberration, and many other devastating cognitive symptoms [1, 2] severely diminishing the quality of life of those affected. Alzheimer’s disease (AD) is the most common form of dementia and is characterized by the presence of extracellular plaques of amyloid-*β* (A*β*) proteins and intracellular fibrils of hyperphosphorylated tau protein (*τP*). As such, it is widely believed that AD is a protein disorder mediated by the brain-wide spreading of A*β* and *τP* aggregates. These toxic protein aggregates then consequently impair the normal functioning of neurons and neural communication. As the disease progresses, the accumulating neuronal damage leads to cognitive decline and clinically-observable changes in brain rhythms as measured by electroencephalography (EEG) and magnetoencephalography (MEG). However, how A*β* and *τP* pathology on the neuronal level affect large-scale brain dynamics—and, by extension, brain function—remains unclear. For example, do AD-induced changes in the brain rhythm spectrum arise from localized neuronal damage or disruptions to the neural connectivity between brain regions?

At the molecular level, the prion-like hypothesis of AD progression proposes that misfolded/modified A*β* and *τP* protein aggregate into soluble, pathological oligomers and spread throughout the brain [3–7]. In particular, AD initiates with A*β* oligomerization spreading diffusively in neocortical regions [8], whereas *τP* spreads—at a later stage—from the entorhinal cortex and continues in a step-wise fashion following axonal pathways between brain regions [8–11]. Studies also suggest that A*β* facilitates the propagation of pathological *τP* during AD progression [11–14]. Although the presence of A*β* is the most reliable diagnostic criteria for AD [15], it is the spreading of pathological *τP* that shows the highest correlation with brain atrophy [16] and cognitive decline [8, 17] leading to the clinical stage of the disease [18, 19].

As A*β* oligomers and *τP* spread throughout the brain, these toxic proteins upset the normal functioning of neuronal cells. Interestingly, recent findings highlight stark differences in the pathology of A*β* and *τP* at the cellular level. On the one hand, toxic A*β* aggregates induce neuronal hyperactivity in rat brain slices and *in vivo* mouse models [20–25] suggested to be caused by activation (deactivation) of excitatory (inhibitory) neurons— shifting the excitatory-inhibitory (E/I) balance towards excitation [26]. On the other hand, *τP* has been linked to axonal damage and loss of white matter integrity in human subjects [27, 28] as well as neuronal hypoactivity in mice models [24, 29– 32]. Interestingly, *τP*-induced hypoactivity predominantly affects excitatory neurons as opposed to inhibitory neurons [33– 35]. Using *in vivo* mouse models, Busche et al. [24] additionally showed that *τP*-induced hypoactivity dominates A*β*-induced hyperactivity in the presence of both A*β* and *τP*. These results inspired Harris et al. [26] to propose the hypothesis that AD follows a biphasic progression, where A*β* oligomers initially induce neuronal hyperactivation which is then later followed by *τP*-induced hypoactivation. However, whether such a hypothesis is compatible with the disparate spreading patterns of A*β* and *τP* as well as AD-related brain rhythm alterations (see below) has not yet been examined. Although the precise mechanisms of neurotoxicity in Alzheimer’s disease is still an open research question, these recent results suggest that *(i)* A*β* oligomers increase excitatory and decrease inhibitory neuronal activity, *(ii) τP* causes axonal damage and decreases the activity of excitatory neurons, and *(iii)* the presence of both *τP* and A*β* decreases neuronal activity.

The malfunctioning of neuronal cells manifests itself on a larger scale in aberrant brain rhythm activity as measured by EEG and MEG. While E/MEG research on AD often reach inconsistent conclusions, some characteristics of the resting-state signal of AD patients have been identified and are reproduced reliably: global frequency-slowing [36–38], power decrease of the dominant rhythm in the alpha frequency band [36– 45], and power increases in both delta [36, 38, 40, 45] as well as theta [37–39, 41–43, 45, 46] frequency bands. Recent findings indicate that oscillatory slowing is prominent in frontal and parietal regions and correlates with subjective cognitive decline [47] and clinical tests for dementia [48]. Moreover, momentary increases in alpha power have been observed for early-stage AD patients [45] and amnesic [47], A*β*-burdened healthy adults [49]. Hyperactivity induced by A*β* and damage-related compensatory mechanisms may be the cause of such early-stage power increases. Indeed, hyperactivity is apparent early in amyloid-precursor overexpressing mice suggesting that A*β* is the primary cause of AD-related hyperactivity [50, 51]. It is worth noting, however, that whether hyperactivity at the neuronal scale causes significant shifts in E/MEG power spectra is still a matter of debate.

Nonetheless, how toxic A*β* and *τP* cause the observed changes to brain rhythm dynamics—and thus, affect cognitive function—remains elusive. And although great progress has been made in understanding the pathological protein spreading and large-scale neuronal dynamics in AD, these processes are often studied in isolation. To date, few attempts have been made at bridging these two processes and shed light on how disease progression affects large-scale brain dynamics and leads to cognitive decline [52].

Here we give insights on how disease-induced changes to brain network structure in AD modulate functionally relevant brain dynamics: Using an integrated computational model that bridges disease progression and neuronal dynamics, we give evidence that local neurodegeneration is the driver behind Alzheimer-like brain rhythm changes. Our computational framework employs a heterodimer model across a connectome to simulate prion-like spreading of two individual proteins—A*β* and *τP*—and monitor the damage imparted on the network. Importantly, we model the effects of A*β* and *τP* separately [53] to account for their distinct effects on different neuron types and axonal integrity. This damage progression then shapes the connectome-level neuronal dynamics as the network deteriorates. We furthermore use our model to test different hypotheses about the neurotoxic effects of proteins. Remarkably, we observe that the key E/MEG brain rhythm characteristics of Alzheimer’s disease are captured. Our simulations reveal a biphasic progression of the disease in which hypoactivity and frequency-slowing are preceded by early-stage hyperactivity. A key insight is that in order to reproduce the slowing of brain rhythms as observed in Alzheimer’s patients, the axonal degradation caused by toxic tau has to be kept small. This finding strongly supports the hypothesis that *local* neurodegeneration is the driver behind Alzheimer-like neuronal dynamics rather than the *global* disruption of connections between brain regions. Using a minimal neuronal population model, we further show that the frequency slowing is caused by decreases in excitatory activity and that stronger synaptic coupling strengths exacerbate the frequency-slowing effect.

## Results

### Formulating a computational framework to investigate Alzheimer’s disease progression

There are three coupled biophysical processes that need to be modeled in order to elucidate the role of neurodegeneration on neuronal dynamics. First, according to both the amyloid cascade and prion-like hypotheses, there is a systematic invasion of the brain by toxic proteins over a typical time scale of 30 years that creates A*β* plaques and *τP* neurofibrillary tangles. The second process is the damage that occurs both at the local regional level and in the connections between regions. The third process is the normal oscillatory dynamics of the neuronal network. Here, we simulate these coupled processes on a connectome, where each node is associated with a brain region. The dynamics can be summarized as follows:

1. Given an initial connection {𝒢}_0_, we simulate the spreading of toxic A*β* and *τP* and compute the damage done by the local concentration of toxic proteins to the connectome {𝒢_*t*_ | *t* ∈ [0, 30]};
2. At given intervals in the sequence *T* = {0, Δ*T*, 2Δ*T*, … }, we simulate large-scale brain dynamics using neural mass models on {𝒢_*t*_ | *t* ∈ *T*} for small time intervals.

Hence, we have to formulate models for three distinct processes: A*β* and *τP* propagation, toxic A*β* and *τP* pathology, and oscillatory brain dynamics. We will now outline our framework which consists of three coupled models; see Materials & Methods section for details on the computational implementation.

### Aβ and τP propagation

We build on the heterodimer model for the prion-like spreading of coupled A*β* and *τP* on an evolving connectome 𝒢_*t*_ with *N* nodes [53]. Initially, the undirected connected connectome 𝒢_0_ is characterized by its weighted adjacency matrix *W* (0) with entries *w*_*ij*_(0) = *n*_*ij*_*/l*_*ij*_ where *n*_*ij*_ is the mean number of axonal fibers connecting node *i* and *j* and *l*_*ij*_ is the mean length of these fibers [52]. The spreading dynamics follows four species 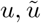 representing healthy/toxic A*β* and 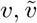 representing healthy/toxic *τP* [53]. The spreading dynamics at node *i* are governed by

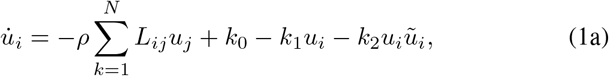

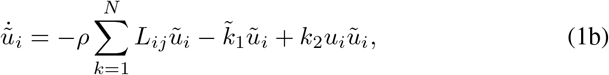

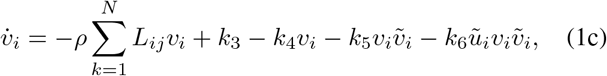

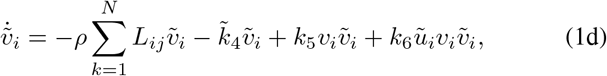

for *i* = 1, …, *N*. Here *ρ* is an effective diffusion constant, the parameters *k*_0_ and *k*_3_ are production rates, 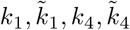, are clearance rates, *k*_2_ and *k*_5_ denote transformations from healthy to pathological proteins, and *k*_6_ describes a synergistic effect between toxic A*β* and toxic *τP* production. The transport between different nodes is characterized by the graph Laplacian

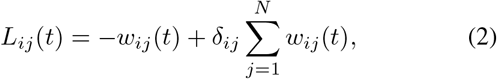

where *δ*_*ij*_ is the Kronecker symbol. This model contains both the conversion from healthy protein to toxic ones and the catalytic effect of *τP* on A*β*.

### Toxic Aβ and tau pathology

In the next step, we proxy the neuronal damage caused by toxic proteins at each node and probe the effect of A*β* and *τP* separately. To this end, we introduce two damage variables 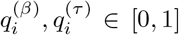 at each node for each protein; here 0 corresponds to no damage and 1 to maximum damage. The damage variables are assumed to follow first-order rate dynamics

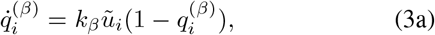

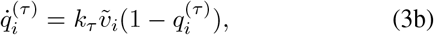

for *i* = 1, …, *N*. Here the parameters *k*_*β*_, *k*_*τ*_ denote the rates at which A*β* and *τP* cause neuronal damage, respectively (see Materials and Methods).

Damage deteriorates both network and nodes. The presence of *τP* affects long-range neural connections (the edges in the connectome). More specifically, if nodes *j* and *k* are initially connected, then the weight of their connection evolves according to

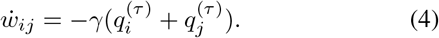

At the level of each individual node, A*β* and *τP* affect the excitatory and inhibitory populations differently. Let *a*_*i*_(*t*) and *b*_*i*_(*t*) denote the activity parameter of the excitatory and inhibitory population at node *i*, respectively. The pathology of A*β* and *τP* is modeled to follow the general trends that: (i) the presence of A*β* increases *a*_*i*_ and decreases *b*_*i*_ (A*β*-induced hyperactivity); (ii) the presence of *τP* decreases *a*_*i*_ (*τP*-induced neurodegeneration); (iii) the presence of both A*β* and *τP* decreases *a*_*i*_ in the long-term (*τP* dominates A*β* effects). A minimal model that expresses these effects is given by

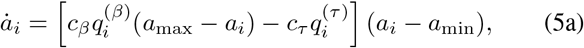

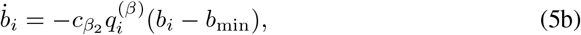

where *a*_*i*_ increases (decreases) when the damage ratio is tilted towards A*β* (tau) pathology and *b*_*i*_ decreases in presence of A*β* pathology.

Together Eqns. (1b–5b) form a system of nine ordinary differential equations that can be solved for given initial conditions. It describes the evolutions of the toxic protein concentration as well as the local and global damage. We integrate these equations over a period of 30 years and probe at regular intervals, every three years, the brain neuronal activity for given fixed parameters at that time *a*_*i*_(*T*), *b*_*i*_(*T*), and *w*_*ij*_(*T*) by a neuronal mass model as shown in Fig. (1d).

**Figure 1:**
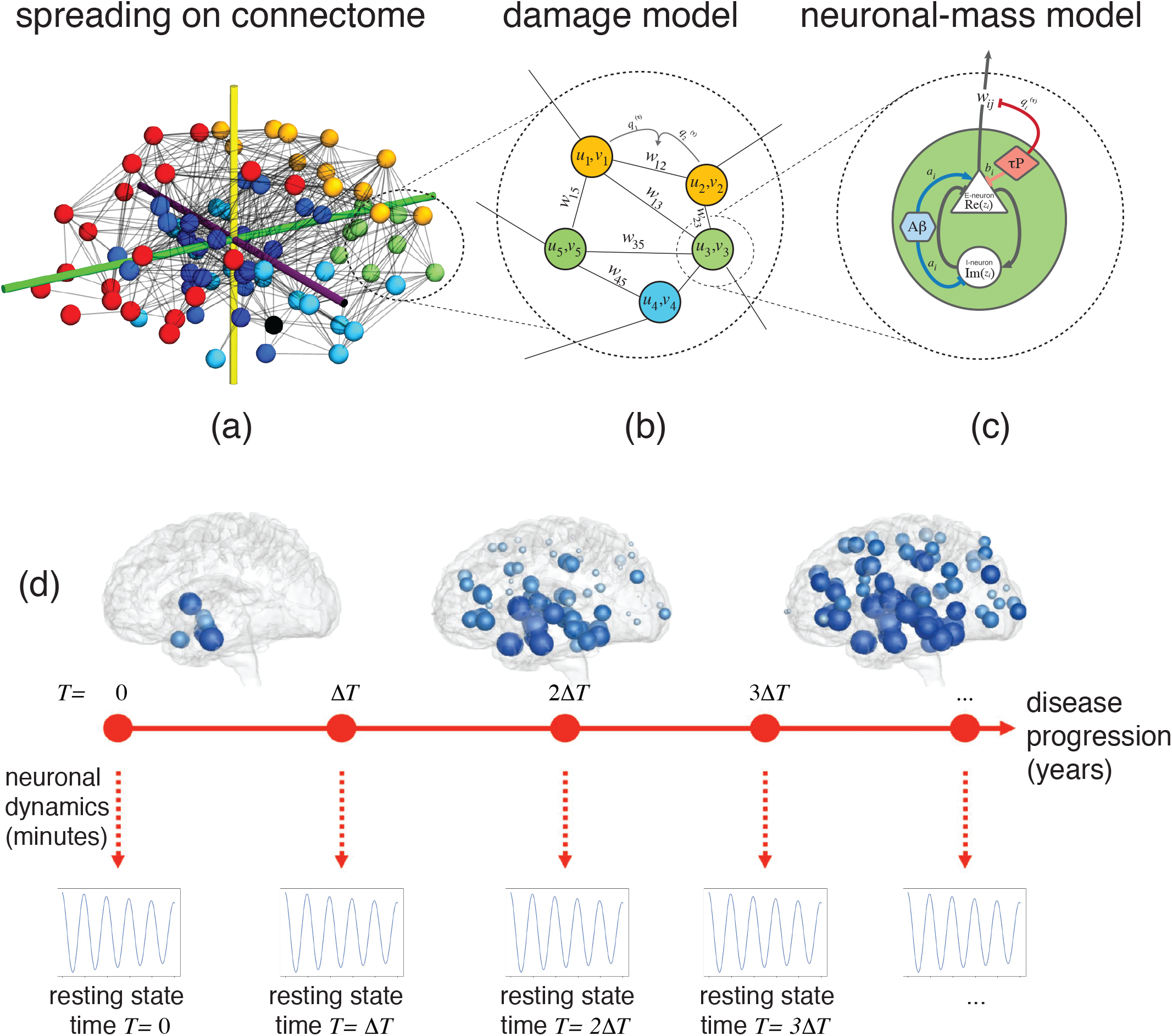
(a) We consider a connectome with *N* nodes, representing brain regions and edges representing axonal connections. (b) On a given node *i*, we define the local concentrations *u*_*i*_ of toxic *Aβ* and *v*_*i*_ of *τP*. Nodes *i* and *j* are connected by an edge with weight *w*_*ij*_(*t*). (c) Each node consists of reciprocally-connected subpopulations of excitatory and inhibitory neurons with their activity parameters *a*_*i*_ and *b*_*i*_. Pathological A*β* disrupts the E/I balance (increases *a*_*i*_, decreases *b*_*i*_), whereas toxic tau damage excitatory neurons (decreases *a*_*i*_) and degrades axonal links (decreases *w*_*ij*_). (d) The dynamics has two different time scales. First, we integrate the spreading and damage model over a period of 30 years. Then, every year, we probe the resting-state dynamics for <1 minute using a neuronal mass model with fixed parameters given by the spreading model.

### Oscillatory brain dynamics

To assess the effect of degeneration on oscillatory brain dynamics, we employ a neuronal mass model on the evolving connectome—a commonly used approach to model whole-brain dynamics. For each node *i* ∈ {1, …, *N*}, let *x*_*i*_ denote the activity of the excitatory population and *y*_*i*_ the activity of the inhibitory populations at the node. Writing the state as a single complex variable *z*_*i*_ = *x*_*i*_ + i *y*_*i*_ (where 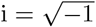), we consider the dynamics

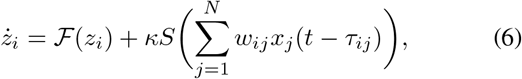

where

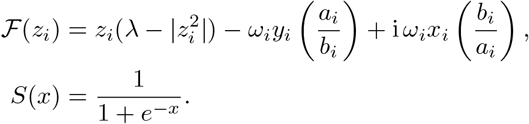

The activity of each node in isolation—determined by ℱ with decay parameter *λ* ∈ ℝ—is governed by the intrinsic frequency *ω*_*i*_ ∈ ℝ close to a Hopf bifurcation with axes corresponding to excitatory and inhibitory activity rescaled by *a*_*i*_, *b*_*i*_ ∈ ℝ, respectively. Coupling between nodes is through the activity of the excitatory activity mediated by the sigmoidal function *S* subject to a time delay of *τ*_*ij*_ ∈ ℝ between nodes *i* and *j*. The neural mass dynamics are much faster compared to the disease evolution (seconds vs years), which justifies our assumption of taking the parameters constant over that time interval.

### Aβ and τP pathology interacts to cause hyperactivity-preceded neurodegeneration

To initialize the disease evolution, we seed toxic *τP* in the entorhinal cortex and toxic A*β* in the precuneus, isthmus cingulate, insula, medial orbitofrontal, and lateral orbitofrontal cortex [54]. The spreading of A*β* and *τP* across different lobes of the connectome is shown in Fig. 2, in which we see a diffusive A*β* spreading followed at a later stage by pathological *τP* spreading from the entorhinal cortices progressively to the neocortex. As the toxic proteins propagate through the brain, we compute how they impart damage to excitatory and inhibitory neuronal populations as well as the deterioration of axonal connections.

**Figure 2:**
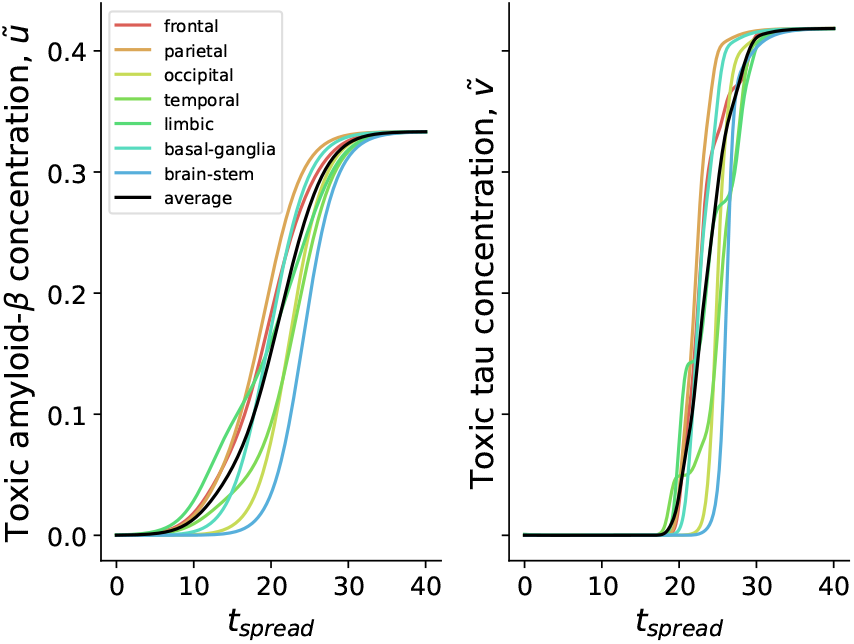
The evolution of toxic amyloid-*β* and *τP* concentration against spreading time. The colored lines correspond to averages of the concentration over different brain lobes, where the black line is the global average.

A*β* and *τP* exhibit distinct pathological characteristics at the neuronal level. Crucially, A*β* and *τP* appear to discriminate between excitatory and inhibitory neurons; this is captured by our primary modeling assumptions is that A*β* and *τP* assert distinctive pathological effects on excitatory and inhibitory nodes. A*β* upsets the E/I balance (increasing excitation, decreasing inhibition) and *τP* damages axonal connections and excitatory neurons (see Eqs. 5a-5b).

Running simulations of A*β* and *τP* spreading across the connectome, we find that the excitatory strength parameter, across all brain regions, decreases after a period of initial hyperexcitability, as can be seen in Fig. 3. Additionally, inhibitory neuronal strength is decreased as excitatory strength peaks. Our simulations predict an initial early-stage E/I imbalance favoring excitation followed by hypoactivation; that is, a biphasic disease progression. However, it still remains to see how the disrupted neuronal parameters affect the large-scale neuronal dynamics themselves. To assess how the disease changes brain dynamics, we simulate neural oscillators on the decaying brain network and compute at regular intervals of Δ*T* = 3 years. Specifically, we calculate the average spectral density across the alpha range (8–12 Hz) over a period of 10 seconds. The evolution of E/MEG alpha spectral density throughout disease progression is shown in Fig. 4. We observe a biphasic disease progression where a period of initial hyperactivity precedes hypoactivity. Regional differences can also be found. While all regions experience hyperactivity, the occipital lobe does so later as compared to other lobes. There is a period in which the occipital lobe is hyper-activated, whereas other brain regions are hypoactivated. The parietal lobe exhibits particularly early hyperactivation, whereas the frontal lobe and limbic system are hypoactivated earlier than other brain regions.

**Figure 3:**
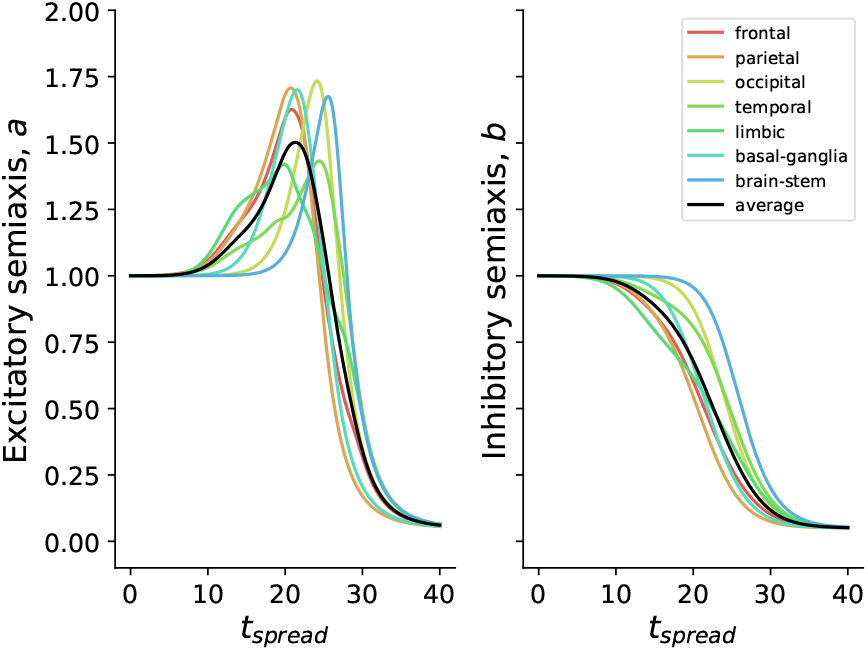
The evolution of neural mass activity parameters *a*_*i*_, *b*_*i*_. The colored lines correspond to averages of *a*_*i*_ and *b*_*i*_ over different brain lobes and the black line is the global average.

**Figure 4:**
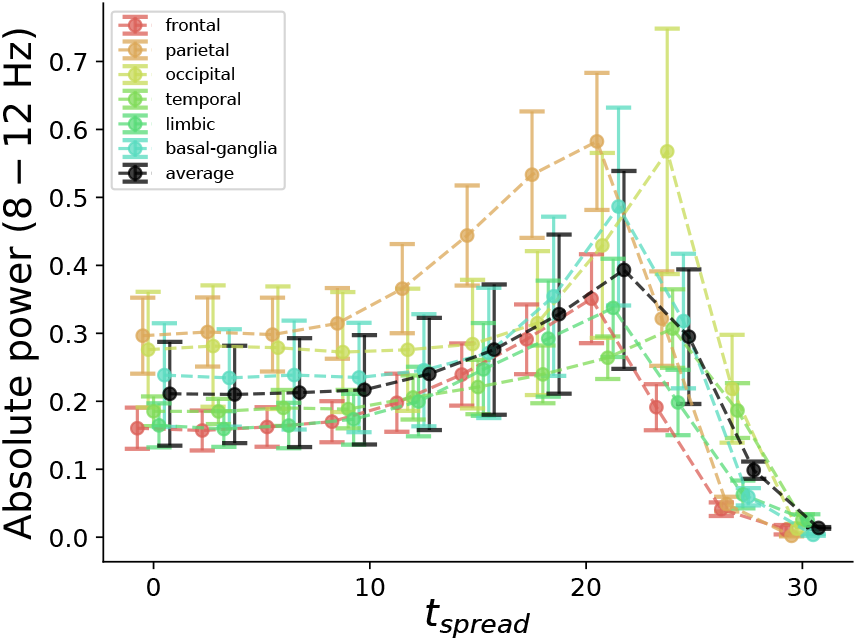
The average spectral density across the alpha range (8-12 Hz) throughout disease progression. The colored lines correspond to averages of the spectral density integral over different brain lobes, where the black line is the global average. The points and error bars are the mean and standard deviation over 10 realizations of normally-distributed intrinsic frequencies.

### Frequency-slowing is induced by local neurodegeneration

One of the most reproducible E/MEG characteristics in Alzheimer’s disease is the slowing of the dominant alpha rhythm. We now show that it is an emerging feature of our model and test whether it is due to local neurodegeneration, axonal degeneration, or both. To this end, we compute the peak frequency (the frequency at which the spectral power reaches its maximum in the 8–12 Hz range) throughout disease progression and vary the effect of axonal degradation induced by *τP* (achieved by changing parameter *γ*, see Methods and Materials). The alpha peaks throughout disease progression are shown in Fig. 5 for three different values of the axonal degradation parameter. For axonal degradation (*γ* = 0.2), we observe a near-monotonic frequency increase which can be easily explained by the gradual decoupling of the neural mass oscillators. Further, we find that oscillatory slowing is only apparent when the axonal degradation caused by *τP* is kept so low that its effect is negligible. We conclude that frequency slowing can be explained by local neurodegeneration alone rather than a change in connectivity. Overall, the different brain regions all follow the same trajectory. However, the parietal and frontal lobes show the most prominent oscillatory slowing. Unlike the frequency peaks, power density curves are mostly unaffected by axonal degradation during disease progression (not shown).

**Figure 5:**
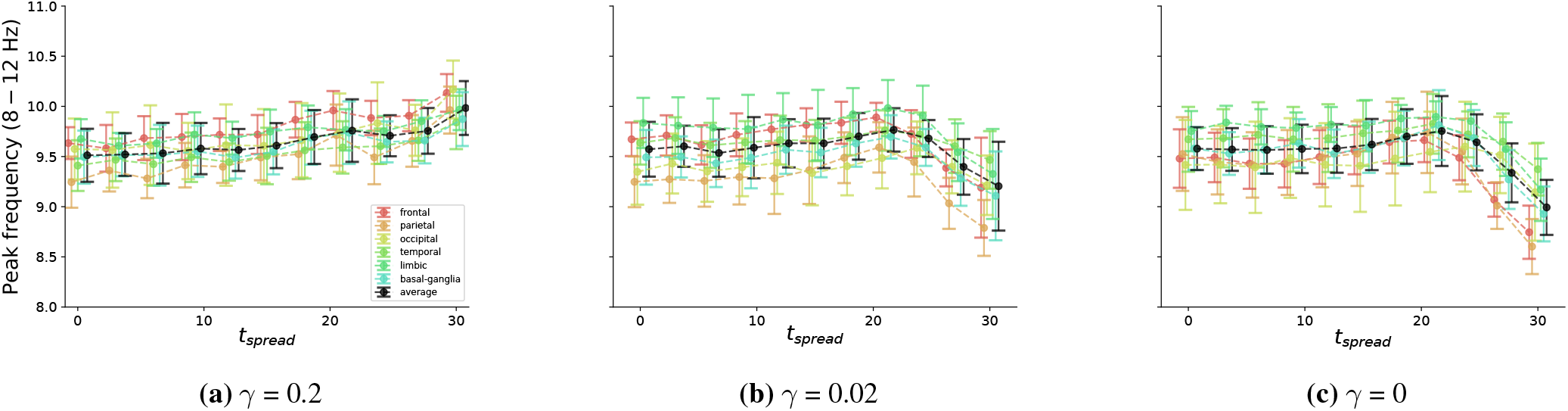
The peak frequency in the alpha band (8 − 12 Hz) throughout spreading time for three different values of *γ* dictating the rate of axonal decay due to *τP* pathology. The colored lines correspond to averages of the peak frequency over different brain lobes, where the black line is the global average. The points and error bars are the mean and standard deviation over 10 realizations of normally-distributed intrinsic frequencies.

### A mechanism for frequency-slowing

The results of the previous section present an interesting phenomenon where frequency-slowing is independent of changes in connectivity and is caused by a decrease in excitatory activity. To understand this effect, we use phase-reduction methods to approximate the dynamics of weakly-coupled oscillators [55]. Consider a single self-coupled node with state *z* = *x* + i *y* that evolves according to

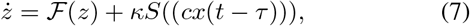

where

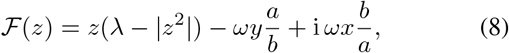

as above, where *κ* is the coupling strength, *a, b* are the excitatory and inhibitory activity parameters, respectively, *ω* is the intrinsic frequency of the node, *c* is the self-coupling link weight, *λ* > 0 is a Hopf-bifurcation parameter, and *S*(*x*) = 1*/*(1+*e*^−*x*^), and *τ* is the time-delay of the self-coupling. By phase reduction, the angle *θ* of points on the limit cycle of the uncoupled system (the “*phase*”) evolves according to

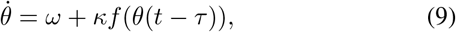

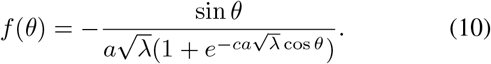

The period *T* for this system is the smallest positive real number such that *θ*(*T*) − *θ*(0) = 2*π* and its frequency of the system, Ω = 2*π/T*. Assuming *τ* small, we obtain

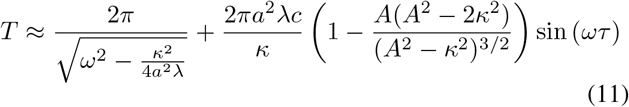

for 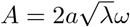 with the approximation *θ*(*t* − *τ*) ≈ *θ* − *ωτ* (see Materials and Methods).

We see that in the absence of delay (*τ* = 0), the frequency decreases as *a* decreases or *κ* increases. Crucially, the decrease in excitatory activity by *a* is sufficient to cause frequency slowing even in an isolated self-coupled node. Hence, local neurodegeneration alone can explain oscillatory slowing. Further, for small *κ* the period can be approximated by

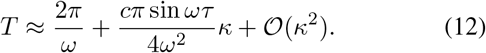

Hence the product of the intrinsic frequency and the delay decides the magnitude and sign of the initial slope of the period *T* against coupling strength *κ*. The effect of the delay on the period is shown in Fig. 6 with a comparison of the original system, the phase-reduced system, and the frequency approximation, showing good agreement. From (12), it is clear that there are parameter regimes where we have frequency slowing or acceleration as shown in Fig. 6.

**Figure 6:**
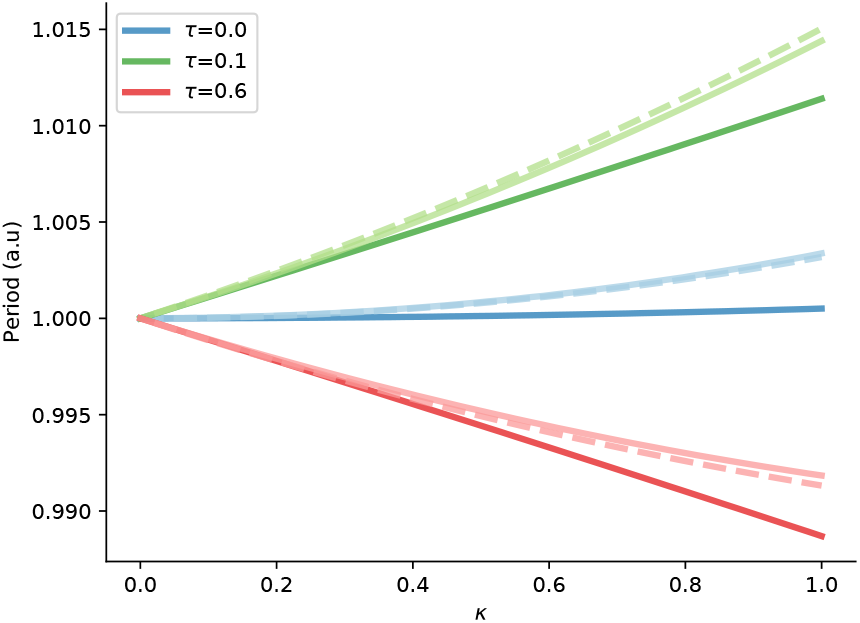
The dependency between period and coupling strength for the self-coupled neural mass model (solid, bold line), its phase-reduced counterpart (solid, light line), and the analytical approximation of the frequency (dashed line). Different colors correspond to different values of the delay *τ*. Other parameters are set as *ω* = 1 Hz, *a* = 1, *b* = 1, *c* = 1, and *λ* = 1.

We conclude that the effect of excitatory activity and coupling strength on frequency is dependent on both the delay and the intrinsic frequency of the corresponding non-coupled system; a phenomenon that can only be captured by introducing delay into the system.

## Discussion

By accounting for the disparate spreading patterns and differential pathology of A*β* and *τP*, we have developed a modeling approach that captures the main characteristics of AD-related brain rhythm alterations. Our approach links the spreading of pathological proteins, their neurotoxic effects, and the temporal evolution of neural dynamics. In doing so, we observe early-stage hyperactivation followed by a decrease and slowing of the dominant alpha rhythm. Moreover, we find that the oscillatory slowing is induced by local neurodegeneration and is independent of damage to interregional connectivity.

In the simulated E/MEG power spectrum, we observe decreases in alpha power following an initial increase in power. It is presently challenging to make comparisons with clinical data, as early-stage hyperactivation in human AD and its effect on E/MEG spectra are yet to be fully explored. Recent studies, however, have found increases in alpha power in healthy A*β*-burdened and amnesic older adults specifically in the parietal region [47, 49], the region which our simulations predict to show the greatest early alpha power increase. Moreover, there is preliminary evidence for hyperactivation in the default-mode network in early-stage AD patients [45]. The spatial heterogeneity in AD-related hyperactivity found in our model could be used as a framework for further experiments in this regard.

Our model robustly predicts the well-documented frequency slowing of the alpha peak. The most affected regions in our simulation are the parietal and frontal lobe, mirroring recent results in subjects showing early signs of dementia [47]. Crucially, we only see this slowing when link degradation is kept small and, as such, implies that axonal damage—though a feature of Alzheimer’s pathology—does not contribute specifically to oscillatory spectral slowing. Further analysis of a minimal neural mass show that frequency slowing is induced by decreases in excitatory neuronal activity (decrease in *a*) and increases in the global coupling strength (increase in *κ*). Frequency slowing has previously been studied in delayed oscillator networks and has been found to be inversely correlated to coupling strengths, average node degree and delay time [56–58]. Interestingly, frequency slowing in our case does not rely on intrinsic frequency heterogeneity, delayed coupling, or network structure. Indeed, even a single self-coupled node can exhibit frequency slowing. All in all, our present work suggests that oscillatory slowing can be explained, parsimoniously and universally, by local node damage rather than an effect due to network connectivity damage.

However, the nature of the change in oscillator frequency becomes more subtle when introducing delay. More specifically, we find that there is an interaction between intrinsic frequencies and delay. For example, decreasing excitation (decreasing *a*) can either impose frequency-slowing or frequency-acceleration dependent on the intrinsic frequency of the system. Hence, delayed neural communication may possibly explain why separate frequency bands are affected differently by A*β* and *τP* pathology. This effect is of particular interest since demyelination has been implied in Alzheimer’s disease [59]. Our model further suggests that changes to excitatory neuronal activity are the primary cause of frequency slowing and not inhibition, which stands in contrast to previous computational studies.

A variety of mechanisms have been proposed to account for the E/MEG spectra slowing as seen in Alzheimer’s patients. These include heterogeneously localized A*β* burden [60], increased inhibition of the thalamus mediated by the basal ganglia [61], and either abnormal increases or decreases in intrathalamus inhibition [62]. Hence, changes to neural circuit excitation have traditionally taken a back seat in explaining deviating E/MEG spectra in AD, despite the observation that restoring excitatory activity levels—in computational models— is the most effective treatment for counteracting AD-related frequency slowing [63] and the relative success of cholinesterase inhibitors—the presently available drugs for AD—which promote signal transmission and have been shown to be able to restore alpha rhythms [64]. Moreover, a recent study found that A*β* and *τP* depositions, respectively, correlate with increases in excitatory and inhibitory time constants of decoupled neural mass models [65] further emphasizing the disparity between A*β* and *τP* pathology. An explanation of this result within our framework is that the presence of *τP* leads to an implicit increase in the excitatory time constant through oscillatory slowing.

While the simplicity of our neural mass model allows for mathematical analysis and a clear interpretation of simulation results, its phenomenological approach yields some limitations. In our approach the intrinsic frequency of each node across the connectome is fixed in the alpha band throughout the brain and, as a phenomenological model, the parameters have limited direct biological interpretation. Thus, in contrast to previous mesoscale computational studies [66], our model lacks a detailed mechanism for the generation of the dominant resting-state alpha rhythm. Moreover, some spatial differentiation of intrinsic frequency may be more realistic such as limiting the generation of alpha-band oscillations to the posterior part of the brain, where they are more typically observed in resting-state E/MEG. Another important consideration is cross-frequency coupling, which has lately been studied in smaller network motifs [67–69] though largely ignored in large-scale connectome models. There is no standard way to incorporate multi-frequency dynamics into brain-wide oscillator networks. One approach is to consider networks of neural masses with multiple dynamical regimes operating at different frequencies [60]. Such a multi-frequency approach would facilitate the validation against experimental data, as clinical studies primarily report relative, not absolute, spectral power changes.

Extensions of our approach, which bridges the biophysical modeling of pathological spreading of A*β* and *τP* to clinical changes in E/MEG spectra, can overcome these limitations. More realistic neural mass models based on mechanistic principles with a multi-frequency repertoire provide a way to get more detailed insights. In this context “next-generation” models [70, 71]—directly linking properties of individual neurons with a mean-field description—are particularly promising. Additionally, a higher-resolution thalamo-cortical circuitry could be embedded into the connectome model, allowing for a more precise description of the posterior alpha rhythm. Our present framework can also be used to study the evolution of functional connectivity. However, if future computational studies are to make precise predictions about the specifics of AD-related changes to neuronal dynamics, the models will have to be tailored by and towards clinical data. This may prove to be a difficult task, as studies investigating E/MEG spectra during Alzheimer usually compare healthy versus unhealthy subjects. That is, to date, there are no longitudinal studies looking at the temporal evolution of E/MEG spectra (or functional networks) during Alzheimer’s disease. Offering a theoretical framework before such clinical studies arrive will hopefully prove helpful. Our present modeling effort also showcases that different aspects of AD can be combined using computational modeling methods, which calls for the incorporation of additional components of the disease such as dysfunctional blood-brain barrier control, demyelination, failings in protein clearance, and inflammatory responses. Demyelination is of particular interest, as it would manifest itself mathematically as a change in delay speed which will differentially affect neural oscillators operating at different frequency bands, as shown by our analysis.

Herein, we have demonstrated that a simple computational approach, testing a recently proposed hypothesis for A*β* and *τP* pathology [26], reproduces hallmark features of AD neuronal dynamics. The spreading patterns and presumed cellular pathology produce a biphasic disease progression, where neuronal hyperactivity is observed before oscillatory slowing and hypoactivity. Furthermore, thanks to the simplicity of our neural mass model, we are able to show that decreases in excitatory neuronal activity induce oscillatory slowing independently of structural changes due to axonal damage. Conclusively, our study suggests that the disparate spreading patterns and neuronal pathology of A*β* and *τP*, together with their distinctive effects on excitatory and inhibitory neurons, may constitute important facets of Alzheimer’s disease and should not be ignored in the computational modeling of the disease.

## Materials and Methods

In our computational framework, we combine the biophysical modeling of the prion-like spreading of A*β* and *τP*, model their damaging effects on the brain network, and then investigate the neuronal dynamics on the decaying network. Here we first detail our computational approach: We report our parametric choices, computational settings, and briefly describe the dynamics of the local damage and neuronal mass model. Details on the prion-spreading model are omitted as they can be found in Thompson et al. (2020). Second, we find the phase-reduced approximation of the single, self-coupled neural mass and derive an analytical approximation for its period of oscillation.

### Prion-like spreading and damage models

The brain network, or connectome, used in our simulations consists of 83 nodes with 579 links and is constructed from tractography of diffusion tensor magnetic resonance images of 418 healthy subjects of the Budapest Reference Connectome v3.0 [72]. The parameters chosen for the simulations in the Results sections are summarized in Table 1. The parameters are composites of units in years and an arbitrary unit of concentration denoted by M. The parameters were chosen close to the exemplary set of parameters for secondary tauopathy as described by Thompson et al. (2020). The parameters *k*_*β*_ and *k*_*τ*_ are chosen so that the evolution of the A*β* and *τP* damage variables closely follow the evolution of toxic A*β* and *τP* concentration. All nodes were initialized identically with respect to healthy protein concentrations, local damage, and excitatory/inhibitory parameters as follows *u*_*i*_(0) = *v*_*i*_(0) = 1 M, 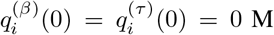, and *a*_*i*_(0) = *b*_*i*_(0) = 1. The initial amount of toxic amyloid-*β* in the entire connectome was 10^−2^ M and was distributed equally among the twelve amyloid-seeded nodes. Likewise, the initial amount of *τP* was also 10^−2^ M distributed between the two tau-seeded nodes. The nonseeded nodes were initialized with zero toxic amyloid-*β* and *τP*. The maximum and minimum values of *a*_*i*_ and the minimum value of *b*_*i*_ were set as *a*_max_ = 1 + *δ, a*_min_ = 1 − *δ*, and *b*_min_ = 1 − *δ*.

**Table 1:** Parameters for the spreading and network pathology simulations.

|  |  |  |
| --- | --- | --- |
| $k_0 = 2 \text{ M years}^{-1}$ | $k_3 = 2 \text{ M years}^{-1}$ | $\gamma = 0.2 \text{ cm years}^{-1}$ |
| $k_1 = 2 \text{ years}^{-1}$ | $k_4 = 2 \text{ years}^{-1}$ | $c_\tau = 1.8 \text{ years}^{-1}$ |
| $k_2 = 2 \text{ M}^{-1} \text{ years}^{-1}$ | $k_5 = 2 \text{ M}^{-1} \text{ years}^{-1}$ | $c_\beta = 0.8 \text{ years}^{-1}$ |
| $\tilde{k}_1 = 1.5 \text{ years}^{-1}$ | $\tilde{k}_4 = 2.66 \text{ years}^{-1}$ | $c_{\beta_2} = 0.4 \text{ years}^{-1}$ |
| $\rho = 1 \cdot 10^{-3} \text{ cm years}^{-1}$ | $k_6 = 12 \text{ M}^{-2} \text{ years}^{-1}$ | $\delta = 0.95 \text{ M}$ |
| $k_\beta = 1 \text{ M}^{-1} \text{ years}^{-1}$ | $k_\tau = 1 \text{ M}^{-1} \text{ years}^{-1}$ | |

### Dynamics of local Aβ and τP pathology

The activity level of the excitatory population is governed by the activity parameter *a* which evolves according to Eq. 5a). Depending on the damage levels, there are typically two stationary points

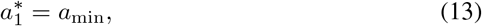

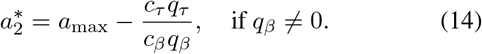

The stability of the two stationary points are interchanged at a transcritical bifurcation (shown schematically in Fig. 7) with respect to the ratio of *τP* to A*β* damage. Letting *R* = *q*_*τ*_ */q*_*β*_, the transcritical bifurcation occurs when *R* crosses *R*_crit_ = (*a*_max_ − *a*_min_)*c*_*β*_*/c*_*τ*_. We have that 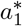 is stable for *R* > *R*_crit_ and 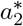 is stable for *R* < *R*_crit_. Note that with toxic A*β* and *τP* present, we have lim_*t*→∞_ *R*_crit_ = 1. Therefore, we pick *R*_crit_ ≤ 1 so that our model is commensurate with experimental findings that *τP* neuronal effects dominate A*β* effects in the long term [24]. By contrast, Eq. 5b which determines the activity parameter *b* of the inhibitory population has only one stable stationary point *b*^*^ = *b*_min_ for *q*_*β*_ > 0.

**Figure 7:**
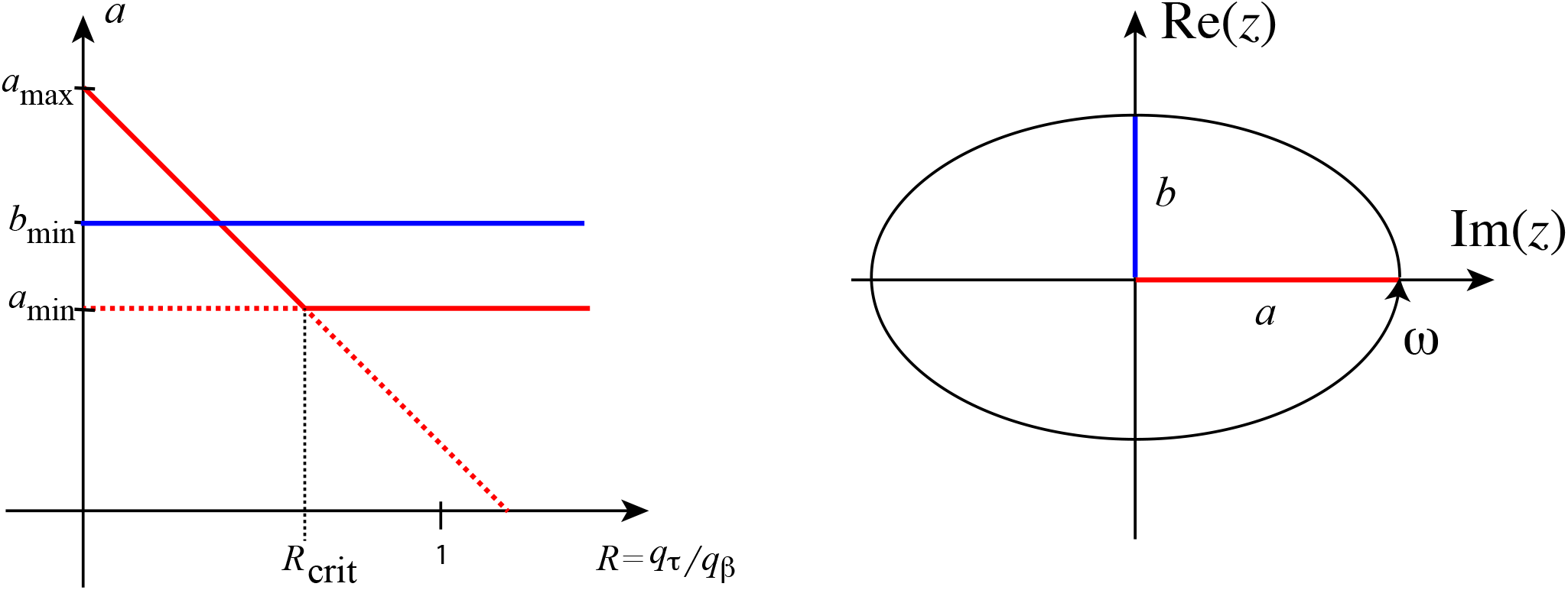
*Left:* An illustration of the transcritical bifurcation observed for the evolution of *a* (red lines). Bold (dashed) lines represent stable (unstable) fixed-points, where we have included the unique stable fixed-point of *b* (blue line). When the damage ratio is tilted towards A*β*-damage (*R* < *R*_crit_), 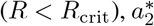 is stable. In contrast, when the damage is tilted towards tau-damage (*R* > *R*_crit_), 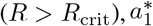 is stable. Note that in our parameterization *a*_min_ = *b*_min_ as opposed to the illustration. *Right:* The stable limit cycle of the (complex) variable *z* = *x* + i *y* in the minimal (without input, *κ* = 0) neural-mass model for λ > 0. The trajectory of the limit cycle follows an origin-centered ellipse with semiaxes (colored red and blue) of length, 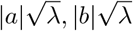, aligned with the real and imaginary axes. The trajectory travels with a constant angular frequency of *ω* with respect to the origin. In the case *λ* < 0, the solutions spiral towards the origin also with a constant angular frequency of *ω*.

### Neural mass model and neuronal dynamics on the connectome

Our neuronal mass model (Eq. 6) generalizes the neural mass model introduced by Goriely et al. (2020), in which node states are modeled by a simplified variant of the Hopf normal form linked with Wilson-Cowan coupling. The intrinsic dynamics of the neuronal mass presented here arise from a scaling of the variables of said model, changing the trajectory of the limit cycle in the complex plane from circular to elliptical. The origin-centered elliptical limit cycle arises from a Hopf bifurcation, having semiaxes with radii 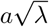 and 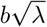 aligned with the *x*-axis (activity of the excitatory population) and *y*-axis (activity level of the inhibitory population), respectively; cf. Figure 7. The Hopf bifurcation occurs at *λ* = 0, where the origin is a stable spiral for *λ* < 0 and the limit cycle becomes attracting for *λ* > 0.

The neural mass parameters chosen for the simulations in the Results sections are summarized in Table 2. We have picked the Hopf bifurcation parameter *λ* close to the point of self-sustained oscillations, so that the oscillations observed in the connectome simulations arise from the input between neighbouring nodes. We compute the delay parameters using an average axonal speed *v*_*ax*_ gathered from the literature [73] by setting *τ*_*ij*_ = *l*_*ij*_*/v*_*ax*_. The delay parameters are further discretized into 40 distinct values to ease computations of the system of delay differential equations. The intrinsic frequencies of the oscillators *ω*_*i*_ are drawn from a normal distribution with mean 10 Hz and standard deviation 1 Hz. Due to the stochasticity invoked by the randomly-drawn frequencies, we repeat the neuronal dynamical simulations 10 times at each time point. The simulation plots in the Results section show averages and standard deviations over these trials. Note however that the brain-stem region is omitted in these plots, as this region consists of a single node rendering averages and standard deviations insubstantial. For each trial, every node variable *z*_*i*_ was initialized at a uniformly-random point inside the complex unit circle. The global coupling strength *κ* was chosen so that sustained oscillations were seen for at least 20 seconds across all nodes in the initial connectome 𝒢_0_. To account for initial transient dynamics, the oscillatory dynamics were run for 20 seconds of which the first 10 seconds were discarded. The remaining 10 seconds of the excitatory time series *x*_*i*_ = Re(*z*_*i*_) are used in the spectral analysis with a sampling frequency of 500 Hz resulting in a frequency resolution of 0.1 Hz. The power spectral density is computed for each node using the *periodogram* function from Scipy’s signal processing module. The absolute alpha power is then computed by integrating the power spectral density from 8 to 12 Hz. The peak frequencies were then taken to be the frequency between 8 and 12 Hz with the largest power spectral density.

**Table 2:**
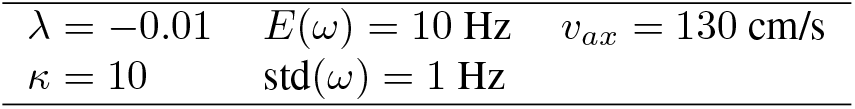
Parameters for the neural mass simulations.

|  |  |  |
| --- | --- | --- |
| $\lambda = -0.01$ | $E(\omega) = 10 \text{ Hz}$ | $v_{ax} = 130 \text{ cm/s}$ |
| $\kappa = 10$ | $\text{std}(\omega) = 1 \text{ Hz}$ | |

### Phase-reduction of self-coupled neural mass

Here we apply methods from phase-reduction theory that enable us to approximate the single self-coupled neural mass by a simpler, one-dimensional system. We will assume a weak self-coupling in order to derive the phase-reduced approximation. First, we will find a phase-description of the uncoupled, single neural mass. A phase-description is a representation of the trajectory of the system through a limit-cycle by a variable *θ* ∈ ℝ*/*2*π* which has a constant derivative. Then, we will approximate the coupling as a function of *θ* by assuming the coupling is weak enough so that the coupled system does not deviate significantly from the uncoupled limit cycle trajectory. Lastly, we will use the phase-reduced system to approximate the period of oscillation under the assumption of weak coupling.

In real vector notation, the state *X* = (*x, y*) of a single, self-coupled neural mass evolves according to

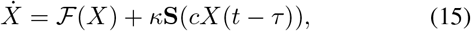

where 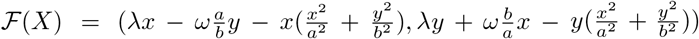, and **S**(*X*) = (*S*(*x*), 0). First, we want to find a phase-description of the uncoupled system. Write the self-coupled system Eq. (15) with *κ* = 0 in scaled polar coordinates (*x* = *ar* cos *ϕ, y* = *br* sin *ϕ*) to obtain

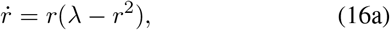

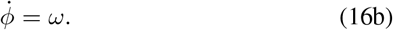

By construction, this system exhibits a Hopf bifurcation at *λ* = 0 with an attractive limit cycle for positive *λ* and an attractive spiral into the origin for nonpositive *λ*. As we are interested in the oscillatory solutions of the system, we will assume *λ* strictly positive. We see that the choice of phase *θ*(*t*) = *ϕ*(*t*) clearly has a constant derivative and is thus a valid phase description for the uncoupled system. As *θ*(*t*) = *ωt*, the limit cycle trajectory is described by

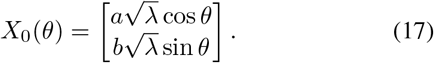

Now note that

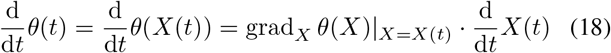

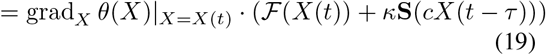

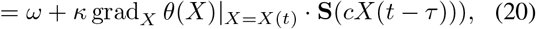

where grad denotes the gradient. To write down the phase reduction, we approximate grad_*X*_ *θ*(*X*)|_*X*=*X*(*t*)_ above by the phase-sensitivity function

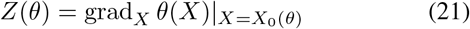

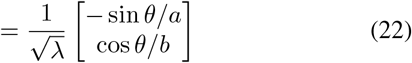

and **S**(*cX*(*t* − *τ*)) by **S**(*cX*_0_(*θ*(*t* − *τ*))). Hence the approximate phase evolution of our self-coupled system is

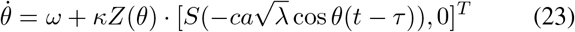

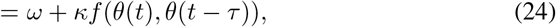

where

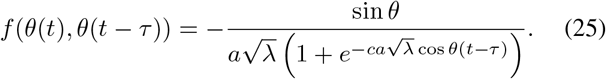

We may use the phase-reduced system to obtain an approximate expression for the period of the system. More precisely, let *θ*(*t*) be a solution to an ODE with at least one stable limit cycle and assume that *θ*(0) = *θ*_0_ is on a stable limit cycle. The *period T of θ*(*t*) is the smallest positive real number such that *θ*(*T*) − *θ*(0) = 2*π*. We will now derive an expression for *T* of the above phase-reduced system (Eq. 9). Starting from the definition of the period, we have

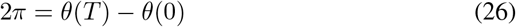

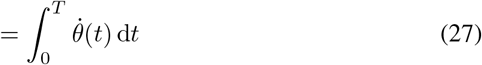

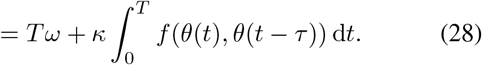

As *κ* is small, we may assume 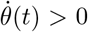 for all *t*. This assumption implies that *θ*(*t*) is a one-to-one function on 0 ≤ *t* < *T* and thereby allows us to use *θ*(*t*) to make an integral substitution above from *t* to *θ*. Hence,

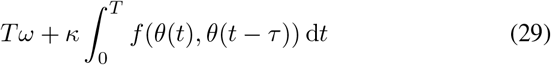

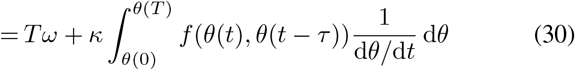

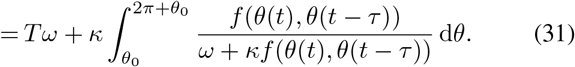

Rearranging in terms of *T* and approximating the delayed phase as if it was free-running, *θ*(*t* − *τ*) ≈ *θ*(*t*) − *ωτ*, we get

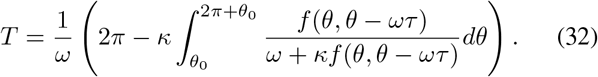

Although we are not able to solve the integral in the above expression analytically, we may solve the integral resulting from a second-order Taylor expansion of the integrand around *c* = 0. Inserting the result of the integral into the above equation leaves us with

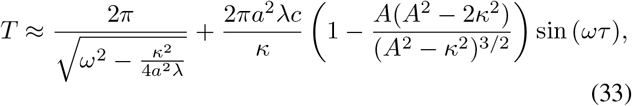

where 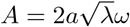, which corresponds to Eq. 11. 10

## Acknowledgments

W.d.H acknowledges support through the NWO ZonMw Memorabel grant (#733050518). C.B. acknowledges support from the Engineering and Physical Sciences Research Council (EPSRC) through the grant EP/T013613/1 as well as the hospitality of the Mathematical Institute of the University of Oxford through an OCIAM Visiting Research Fellowship. The support for A.G. by the Engineering and Physical Sciences Research Council of Great Britain under research grant EP/R020205/1 is gratefully acknowledged.

